# IL-1β-mediated Immunometabolic Adaptation in Corneal Epithelial Cells

**DOI:** 10.1101/2024.09.08.611874

**Authors:** Jose Marcos Sanches, Rajalakshmy Ayilam Ramachandra, Natalia Mussi, Hamid Baniasadi, Danielle M. Robertson

## Abstract

**Purpose:** As an external mucosal surface, the corneal epithelium is subject to a barrage of stressors that are known to trigger inflammation. IL-1β, a master regulator of inflammation, is secreted into the preocular tear film by ocular surface epithelial cells and infiltrating immune cells. While increased levels of IL-1β have been associated with corneal disease, the effects of IL-1β on mitochondrial function in corneal epithelial cells (CECs) is unknown.

**Methods:** To investigate the effects of IL-1β on mitochondrial function, telomerase immortalized human CECs were cultured in either 50 ng/mL or 100 ng/mL IL-1β for short term (24 hours) or prolonged (72 hours) time periods. Cells were assessed for ROS, inflammatory cytokine production, mitochondrial polarization and ultrastructure, mitophagy, and changes in the metabolite composition. Lipid drops were examined using light and fluorescent microscopy.

**Results:** Short term exposure to IL-1β triggered an increase in IL-8 and ROS levels that corresponded to a reduction in mitochondrial membrane potential. Long term exposure also showed increased levels of IL-8 and IL-6 and further increased ROS. After long term exposure however, there was a paradoxical increase in mitochondrial membrane potential that was associated an increase in spare respiratory capacity and mitochondrial hyperfusion. Metabolomics confirmed an upregulation of the pentose phosphate pathway and the TCA cycle. Fumarate was also increased, suggesting an increase in flux through complex II. Changes in lipid metabolism included an upregulation in cardiolipin and *de novo* triacylglyceride biosynthesis, along with increasing numbers of lipid droplets.

**Conclusion:** Prolonged exposure to IL-1β induces metabolic rewiring in CECs that results in an increase in spare respiratory capacity. These findings suggest that the corneal epithelium is able to adapt to certain levels of chronic inflammation and may have important implications in our understanding of immune tone and cellular stress responses in ocular surface epithelia.

## Introduction

The corneal epithelium is a stratified epithelial sheet that together with the conjunctival epithelium forms the anteromost surface of the eye. Like other mucosal surfaces, the corneal epithelium is constantly bombarded by physiological stress induced from eyelid blinking and hypoxia during overnight eye closure, and pathological stress due to increases in tear film osmolarity, desiccation, pollutants, and glucose.^1–18^ A consequence of pathological stress is the induction of inflammation and subsequent surface epithelial cell damage. Interleukin-1β (IL-1β) is a pro-inflammatory cytokine that functions as a master regulator of inflammation.^19^ In response to stress, corneal and conjunctival epithelial cells along with infiltrating immune cells, secrete IL-1β into the precorneal tear film.^20^ In the wounded corneal epithelium, an increase in tear levels of IL-1β is associated with delayed wound healing.^21^ Increased levels of IL-1β have been demonstrated in humans and animal models of dry eye disease.^20,22^ In autoimmune-regulator (aire)-deficient mice, a model for Sjögren’s Syndrome, IL-1ý has been linked with the onset of squamous metaplasia.^23^ At the cellular level, IL-1β exerts pleiotropic effects on the corneal epithelium. In cultured corneal epithelial cells, exposure to IL-1β has been shown to disrupt barrier function through activation of NF-_K_B.^24^ In Statens Seruminstitut Rabbit Corneal Cells, IL-1β stimulated cell migration through the upregulation of matrix metalloproteinase 9.^25^ Using a mouse cauterization model, IL-1β has also been shown to play a pivotal role in the induction of corneal neovascularization.^26^

Mitochondria are dynamic organelles that orchestrate a diverse array of cellular stress responses.^27^ Despite being well known for their role in ATP production through oxidative phosphorylation, mitochondria also regulate cytosolic calcium levels and serve as a primary source of reactive oxygen species (ROS). As major signaling hubs, mitochondria mediate many vital metabolic processes including amino acid catabolism and anabolism, the biosynthesis of macromolecules, pyrimidine and purine metabolism, and the regulation of apoptosis. More recently, mitochondria have been implicated in mediating inflammation and cell autonomous immunity. The mechanisms by which this occurs are varied and include the release of mtDNA, mtRNA, ROS, and IL-1β.^28,29^ In order to maintain cell and tissue homeostasis during stress, mitochondria progress through one of three general pathways. These include an increase in mitochondrial fusion to dilute out damaged mitochondria and maintain cellular bioenergetics, mitophagy to remove damaged mitochondria via lysosomal degradation, or when severe, apoptosis.^30^ Of these, mitophagy is a critical mitochondrial quality control mechanism.^31^

We have previously reported that hyperosmolar and hyperglycemic stress impairs mitochondrial metabolism in corneal and conjunctival epithelial cells.^5–7,13^ In the presence of a mild hyperosmolar stress, there is an increase in oxidative phosphorylation that becomes suppressed at higher levels.^7^ This suppression is associated with a loss in mitochondrial polarization and an increase in PTEN-induced kinase 1 (PINK1)-mediated mitophagy.^6^ In the context of diabetes, we further demonstrated that chronic long term exposure to hyperglycemia is required to induce mitochondrial dysfunction, whereas short term exposure is sufficient to trigger a loss in spare respiratory capacity.^13^ Spare respiratory capacity is an important indicator of cell health.^32^ The loss of spare respiratory capacity creates the potential for a bioenergetic crisis in the presence of additional stress or trauma. Such is the case for diabetes, where the hyperglycemia induced loss of spare respiratory capacity may contribute to a delay in wound healing.^13^

In the present study, we used the pro-inflammatory cytokine IL-1β to investigate the effects of inflammation on mitochondrial and metabolic homeostasis in corneal epithelial cells. While IL-1β triggered an initial loss of mitochondrial membrane depolarization that was associated with an increase in ROS production, prolonged culture induced metabolic adaptation characterized by the restoration of membrane polarization and an increase in spare respiratory capacity. Taken together, these findings suggest that the corneal epithelium is able to adapt to certain levels of chronic inflammation and may have important implications in our understanding of immune tone of the ocular surface.

## Material and methods

### 2.1 Cell culture

A human telomerized corneal epithelial (hTCEpi) cell line previously developed and characterized by our laboratory, was used for these studies.[41] hTCEpi cells were maintained in serum-free keratinocyte growth medium containing 0.06 mM calcium and 6 mM glucose (VWR, Radnor, PA). Calcium levels were increased to 0.15 mM calcium and media was further supplemented with bovine pituitary extract (0.0004 mL/mL), recombinant human epidermal growth factor (0.125 ng/mL), recombinant human insulin (5 µg/mL), hydrocortisone (0.33 µg/mL), epinephrine (0.39 µg/mL), and human holo-transferrin (10 µg/mL, PromoCell; VWR, Radnor, PA). Cultures were maintained at 37^°^C with 5% CO_2_. For all experiments, cells were seeded into culture dishes and allowed to adhere overnight. Media was then changed to basal media without growth factor supplements. To simulate inflammation, cells were incubated with the pro-inflammatory cytokine IL-1β (PeproTech, Rocky Hill, NJ) at a concentration of 50 ng/mL or 100 ng/mL for 24 hours (short term exposure) or 72 hours (prolonged exposure).

### 2.2 Quantification of cell number

To quantify the number of hTCEpi cells at each time point, hTCEpi cells were seeded at a density of 1 × 10^5^ cells per well in a 6-well culture plate. Cells were cultured in basal media with or without 50 ng/mL or 100 ng/mL of IL-1β. Cells were imaged at the indicated time points using a Celigo Imaging Cytometer (Nexcelom Biosciences, Lawrence, MA) and counted using the manufacturer provided software. Each experiment was performed in triplicate and repeated a minimum of two additional times.

### 2.3 Lactate dehydrogenase assay

Lactate dehydrogenase (LDH) release was quantified using the Pierce LDH Cytotoxicity Assay Kit (Thermo Fisher Scientific, Waltham, MA), according to the manufacturer’s instructions. hTCEpi cells were seeded at a concentration of 1 × 10^4^ cells per well in a 96 well culture plate. After overnight attachment, media was removed and cells were cultured in basal media with or without 50 ng/mL or 100 ng/mL of IL-1β. Supernatants were then removed and tested for lactate dehydrogenase according to manufacturer instructions. Briefly, supernatants were added to 25 μl of buffer in each well of a 96 well plate. Substrate was added and samples were incubated at 37°C for 15 minutes. This was followed by the addition of the chromogenic agent for an additional 15 minutes. Finally, the alkali reagent was added and cells were allowed to stand at room temperature for 5 minutes. Absorbance was measured at 490 nm using a BioTek Synergy 2 Multi-Mode Microplate Reader (Thermo Fisher, Waltham, MA). All experiments were performed in four replicates and repeated two additional times.

### 2.4 Intracellular ROS assay

The effect of IL-1β on intracellular reactive oxygen species (ROS) levels was measured using CellROX™ Green Reagent (Thermo Fisher Scientific, Waltham, MA). hTCEpi cells were seeded onto coverslip bottom MatTek culture dishes (MatTek Corporation, Ashland, MA) at a density of 2 x 10^5^ cells per dish. Cells were then treated with IL-1β for 24 or 72 hours. Following treatment with IL-1β, cells were incubated with 5 μM of CellROX® Reagent for 30 minutes at 37°C. Media was then removed and the cells were washed with phosphate buffered saline (PBS), fixed with 4% formaldehyde for 15 minutes at room temperature, and mounted with Fluoroshield™ containing 4′,6-diamidino-2-phenylindole (DAPI, Sigma Aldrich, Saint Louis, MO, USA). Cells were imaged on a Leica SP8 laser scanning confocal microscope (Leica Microsystems, Heidelberg, Germany) using a 63× oil objective. Excitation lasers for DAPI and CellRox were 405 nm and 488 nm, respectively. Cells were sequentially scanned to avoid spectral crosstalk.

### 2.5. JC-1 staining

To determine the effects of IL-1β on mitochondrial polarization, 2 x 10^5^ hTCEpi cells were cultured on coverslip bottom MatTek dishes (MatTek Corporation, Ashland, MA). After treatment with IL-1β for the indicated time period, cells were labelled with 10 µg/mL of tetraethyl-benzimidazolylcarbocyanine iodide (JC-1) dye (Invitrogen/Molecular Probes, Eugene, OR). For a positive control for depolarization, cells were treated with the oxidative phosphorylation uncoupler, carbonyl cyanide m-chlorophenyl hydrazine (CCCP, 50 nM), for 10 minutes at 37°C. Cells were then washed with PBS. JC-1 staining was visualized using a Leica SP8 laser scanning confocal microscope (Leica Microsystems, Heidelberg, Germany) with a 63 × oil objective. The microscope is equipped with an environmental chamber to maintain cells at 5% CO2 and 37°C during imaging. JC-1 monomers were scanned using a 488 nm excitation laser and JC-1 aggregates were scanned using a 561 nm excitation laser. Sequential scanning was performed to prevent spectral crosstalk. Imaging experiments were repeated an additional two times.

### 2.6 Real-time metabolic studies

Real-time measurements of cellular oxygen consumption rate (OCR) and extracellular acidification rate (ECAR) were performed using a Seahorse Metabolic Analyzer XFp (Agilent Technologies, Santa Clara, CA). hTCEpi cells were seeded in Seahorse XFp mini plates at a concentration of 32,000 cells per well and treated with 50 ng/mL or 100 ng/mL IL-1β for 72 hours. Basal media without IL-1β served as a non-treated control. Prior to initiating measurements, cells were first incubated for 1 hour at 37°C in Seahorse XF Dulbecco’s modified Eagle’s medium (DMEM) containing 5 mM 4-(2-hydroxyethyl)-1-pipera-zineethanesulfonic acid (HEPES, pH 7.4) and supplemented with 1 mM pyruvate, 2 mM glutamine, and 10 mM glucose in a non-CO_2_ incubator. OCR and ECAR were analyzed using a Seahorse XFp Cell Mito Stress Test kit (Agilent Technologies, Santa Clara, CA, USA). All assays were performed according to manufacturer instructions. For the XFp Cell Mito Stress assay, the first injection was 10 µM oligomycin to inhibit ATP synthase, followed by one injection of 5 µM carbonyl cyanide 4-(trifluoromethoxy) phenylhydrazone (FCCP) and a final injection of 5 µM of rotenone/antimycin A (R/A). FCCP is an uncoupler of oxidative phosphorylation and R/A are inhibitors of complex I and III, respectively. Treatment with R/A allows for complete inhibition of mitochondrial respiration and determination of non-mitochondrial respiration. Mean OCR and ECAR values were calculated from the measurement obtained at the third time point prior to the addition of any test compounds. The ratio for OCR/ECAR was also calculated for each experiment. Basal respiration was calculated as OCR minus non-mitochondrial oxygen consumption. The proton leak and ATP-linked respiration were also evaluated. Spare respiratory capacity was calculated by subtracting basal respiration from FCCP-stimulated OCR (maximal respiration). Data were analyzed using the manufacturer-provided Wave software version 2.3.0. Samples were plated in 6 replicate wells. All outcome measures were normalized to total cell number using a Celigo imaging cytometer (Nexcelom Bioscience, Lawrence, MA). Each experiment was repeated a minimum of three additional times.

### 2.7 Enzyme-Linked Immunoassay (ELISA)

Human IL-8/CXCL8, IL-6 and IL-10 Quantikine enzyme-linked immunoassays (R&D systems, Minneapolis, MN) were used to analyze secreted cytokine levels in conditioned media. Cells were seeded in six-well plates and allowed to adhere overnight. Media was removed and cells were then cultured in basal media with or without 50 ng/mL or 100 ng/mL of IL-1β and incubated for 24 or 72 hours. For the 24 hour treatment, 7.5 x 10^5^ hTCEpi cells were seeded in six well plates. For the 72 hour time treatment, 5 x 10^5^ cells were in six well plates. Media were collected and concentrated using protein concentrators containing a polyethersulfone membrane (3K MWCO; Millipore, Burlington, MA). Protein concentration was measured using a Qubit 3.0 Fluorometer (Thermo Fisher, Waltham, MA). ELISAs were performed according to manufacturer recommendations. All experiments were performed in triplicate and repeated a minimum of two additional times. Adherent cells were used for immunoblotting, as described below.

### 2.8 Immunoblotting

For immunoblotting of whole cell lysates, a lysis buffer containing 50 mM Tris-HCl pH 7.5, 150 mM NaCl, 1% Triton X-100, 1 mM EDTA, and a protease and phosphatase inhibitor cocktail (Thermo Fisher, Rockford, IL) was used to directly lyse adherent epithelial cells in 6-well culture plates. Samples were kept on ice and vortexed intermittently for 30 minutes, followed by centrifugation for 5 minutes at 12,000 rpm at 4°C (Bio-Rad, Hercules, CA). After centrifugation, the supernatants were collected and protein concentration was quantified using a Qubit 3.0 Fluorometer (Thermo Fisher, Rockford, IL, USA). A 2× sample buffer pH 6.8 containing 65.8 mM Tris-HCl, 26.3% (**w**/**v**) glycerol, 2.1% SDS, 5.0% β-mercaptoethanol and 0.01% bromophenol blue was added to the sample, which was then boiled for five minutes (Bio-Rad, Hercules, CA, USA). Next, boiled samples were electrophoresed through a 4–15% precast linear gradient polyacrylamide gel (Bio-Rad, Hercules, CA, USA). Once electrophoresis was complete, gels were transferred onto a polyvinyl difluoride (PVDF) membrane (Millipore, Temecula, CA, USA). Membranes were then blocked with 5% non-fat milk for one hour at room temperature (Bio-Rad, Hercules, CA, USA). After blocking, membranes were washed three times for five minutes with PBS. Membranes were then incubated overnight in primary antibody at 4°C. Membranes were again washed three times for five minutes with PBS. The primary antibodies that were used included anti-rabbit PINK1, anti-rabbit BNIP3L/NIX, and anti-rabbit vinculin (Cell Signaling, Danvers, MA). Membranes were then incubated with an anti-rabbit secondary antibody at room temperature for one hour (Santa Cruz, CA). An Amersham Imager 600 (Amersham Biosciences, Piscataway, NJ) was used to image membranes. Antibodies were detected using ECL Plus Prime Detection Reagent (Amersham Biosciences, Piscataway, NJ, USA). Vinculin or β-actin were used as loading controls for normalization. ImageQuant TL Toolbox v8.1 software was used to analyze and quantify relative expression of the indicated proteins (arbitrary units, a.u.).

### 2.9 Transmission electron microscopy

To examine mitochondrial ultrastructure, transmission electron microscopy was performed at the UT Southwestern Electron Microscopy core facility. hTCEpi cells were seeded onto 35 mm coverslip bottom dishes and allowed to adhere overnight. Cells were then treated with or without 100 ng/mL IL-1β in basal media for 72 hours. At the end of the treatment period, cells were fixed for 15 minutes at room temperature using 2.5% glutaraldehyde/0.1 M cacodylate buffer buffer (pH 7.4). Next, cells were washed with 0.1 M sodium cacodylate buffer and then incubated with 1% osmium tetroxide + 0.8% K3[Fe(CN6)] in 0.1 M sodium cacodylate buffer for 1 hour at room temperature. Cells were then stained overnight with 2% aqueous uranyl acetate. The following day, cells were dehydrated with ethanol, infiltrated with Embed-812 resin, and then polymerized at 60°C overnight. Sections were cut using a diamond knife (Diatome) on a Leica Ultracut UCT (7) ultramicrotome (Leica Microsystems, Wetzlar, GE). Stained sections were then placed on copper grids and the samples were post-stained with aqueous 2% uranyl acetate and lead citrate. Images were captured using a JEOL 1400 Plus (JEOL) transmission electron microscope equipped with a LaB6 cathode.

### 2.10 Lipid accumulation

To determine the effects of IL-1β on mitochondrial polarization, hTCEpi cells were seeded at a density of 2 x 10^5^ per dish on coverslip bottom MatTek dishes (MatTek Corporation, Ashland, MA). After treatment with 100 ng/mL IL-1β for 72 hours, cells were then labelled with Oil Red O (StatLab, American MasterTech, Lodi, CA, USA) or HCS LipidTox Neutral Lipid Stain (Invitroge, Eugene, OR, USA) following manufacturer guidelines. Cells were visualized using a Leica DMI3000 B microscope (Leica Microsystems, Heidelberg, Germany) with a 63 × oil objective. For LipidTox staining, using a filter cube for Texas Red. Imaging experiments were repeated for at least three times.

### 2.11 Metabolomics analysis

For metabolomic studies, 1.5 x 10^6^ hTCEpi cells were seeded in 10 cm dishes and treated with 100 ng/mL IL-1β for 72 hours. Following treatment, media was removed. A solution consisting of 80% methanol (HPLC grade) with 0.2 µM of amino acids heavy standard (-80^°^C) was then added to the adherent cells and incubated for 20 minutes at -80°C. Cell extracts were collected using a cell scraper and centrifuged at 14,000g for 5 minutes at 4^°^C. Supernatants containing the metabolites were transferred to a new tube on dry ice. The pellet was then washed with 500 µl of 80% methanol solution followed by centrifugation at 14,000g. The samples were filtered and stored in glass vials at -80^°^C until analysis. LC-MS/MS mass spectrometric analyses were performed at the UT Southwestern Metabolomics Core facility using a Sciex “QTRAP 6500+” mass spectrometer equipped with an ESI ion spray source. The ESI source was used in both positive and negative ion modes. The ion spray needle voltages used for MRM positive and negative polarity modes were set at 4800 V and −4500 V, respectively. The mass spectrometer was coupled to a Shimadzu HPLC (Nexera X2 LC-30AD). The system is controlled by Analyst 1.7.2 software. Chromatography was performed using a SeQuant® ZIC®-pHILIC 5 μm polymeric 150 × 2.1 mm PEEK coated HPLC column, (MilliporeSigma). The column temperature, sample injection volume, and flow rate were set to 45°C, 5 μL, and 0.15 mL/min, respectively. The HPLC conditions were as follows: Solvent A: 20 mM ammonium carbonate including 0.1% ammonium hydroxide and 5µM of medronic acid. Solvent B: acetonitrile. Gradient condition was 0 min: 80% B, 20 min: 20% B, 20.5 min 80% B, 34 min: 80% B. The total run time was 34 minutes. Data was processed by SCIEX MultiQuant 3.0.3 software with relative quantification based on the peak area of each metabolite. For LC-MS data, before statistical analysis, the peak area of each feature was normalized against the internal standard. Afterward, statistical analysis was performed using the MetaboAnalyst 4.0 web platform (McGill University, Montreal, Canada). Raw data were normalized and range-scaled before performing statistics. Only those features, such as ions with unique m/z and retention times that fulfilled the criteria of adjusted p-value/false discovery rate (FDR) < 0.05 were considered significant. Adjusted p-values/FDRs and hierarchical analysis were used to construct heat maps.

### 2.12 Statistical analysis

All data are presented as mean ± standard deviation. For the comparison between two groups with a normal distribution, a student’s t-test was performed. For comparisons between more than two groups, a one-way ANOVA was used with an appropriate post hoc multiple comparison test. Statistical significance was set at p<0.05.

## Results

### IL-1β induces pro-inflammatory cytokine release and ROS in hTCEpi cells independent of mitochondrial membrane potential

To test the effects of IL-1β on hTCEpi cells, cells were treated for 24 (short term) or 72 (prolonged) hours. Secretion of IL-8, IL-6, and IL-10 was quantified using ELISA. After short term exposure to IL-1β, IL-8 secretion was increased for both 50 ng/mL IL-1β (P=0.022) and 100 ng/mL IL-1β (P=0.005, Figure 1A). There was no increase in IL-6 or IL-10 at either concentration (Figure 1, B-C). After prolonged expousre, IL-8 remained high (P<0.0001, Figure 1D). IL-6 was also increased by treatment with 100 ng/mL IL-1β (P=0.037, Figure 1E). There was no effect on IL-10 (Figure 1F). We next quantified the effect of IL-1β on intracellular ROS production. Short term exposure showed increasing levels of IL-1β that corresponded to increasing levels of ROS (Figure 1G). Prolonged exposure to IL-1β also showed a small increase in ROS in control cells. Since cells were cultured in basal media over the 72 hour period, this reflects the increase in ROS in response to growth factor withdrawal. Compared to the control, stimulation with IL-1β further increased ROS in a concentration dependent manner. To confirm that the increase in ROS was not associated with any measurable level of cytotoxicity, cell viability was determined using LDH assays and cell counts. Importantly, there were no changes in either parameter at any of the time points and concentrations tested (Figure 2, A-D).

**Figure 1:**
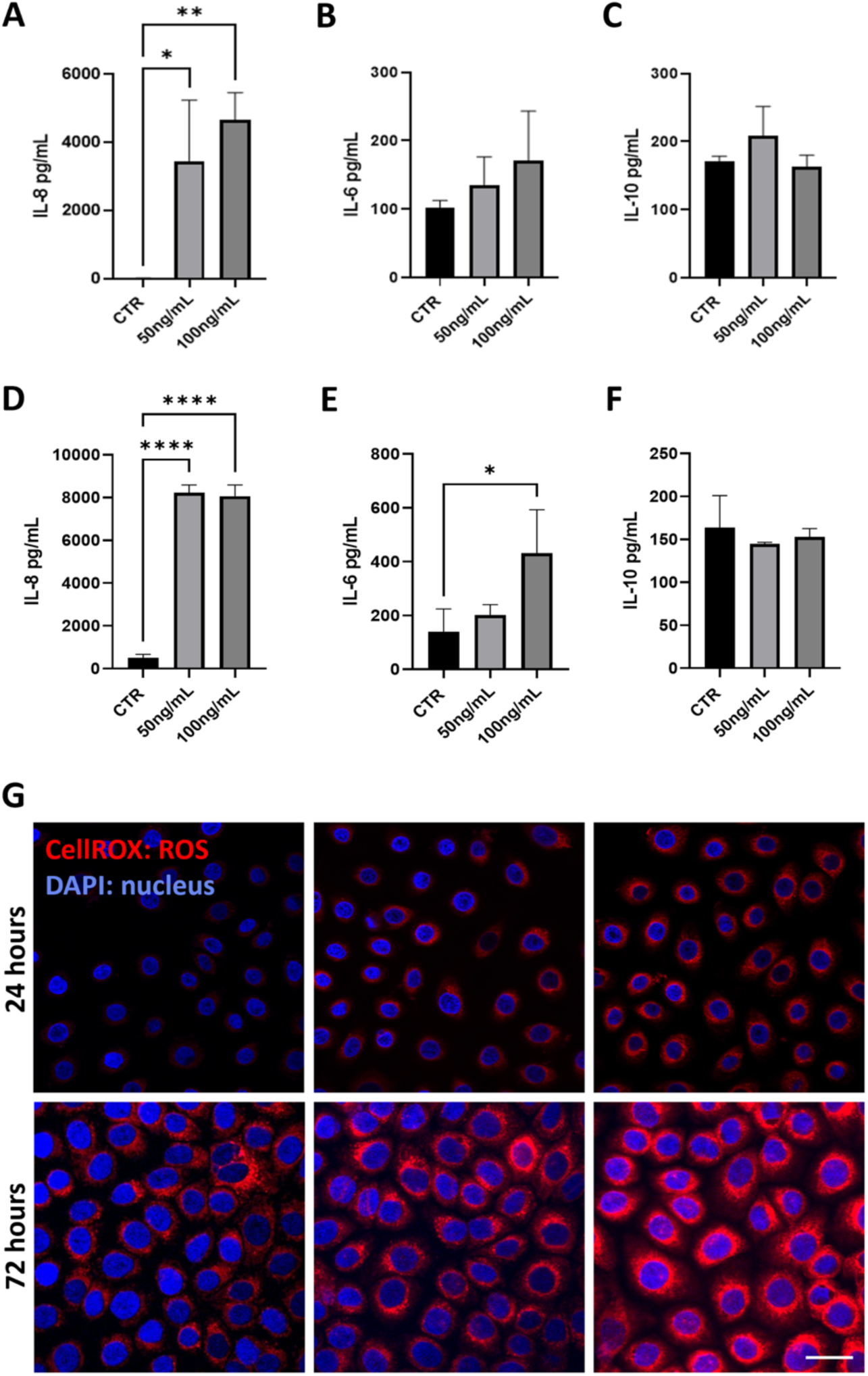
IL-1β-induced inflammation increases pro-inflammatory cytokines release and ROS in corneal epithelial cells. (A-F) ELISA was used to quantify secretion of IL-8, IL-6, and I-10. A: IL-8 release from hTCEpi cells was significanctly increased after 24 hours of treatment with IL-1β in a dose-dependent manner (p=0.023 and p=0.006 for 50 and 100 ng/mL, respectively). (B) While IL-6 levels trended upward, these changes were not signficant. (C) IL-1β did not alter IL-10 release. (D) After 72 hours of treatment with IL-1β, IL-8 was significantly increased at both concentrations (p<0.0001). (E) IL-6 was increased at 72 hours after treatment with 100 ng/ml IL-1β (P=0.04). (F) IL-1β did not alter IL-10 release at 72 hours. (G) CellROX staining for ROS. ROS shown in red, nuclei labeled with DAPI (blue). Treatment with 50 ng/mL and 100 ng/mL of IL-1β showed an increase in ROS levels at 24 hours. ROS was further increased at 72 hours. Scale bar: 20 μm. All data are epressed as mean ± standard deviation of one representative experiment, N=3. One-way ANOVA with a Tukey’s post hoc multiple comparisions test. Images representative of 3 repeated experiments.

**Figure 2:**
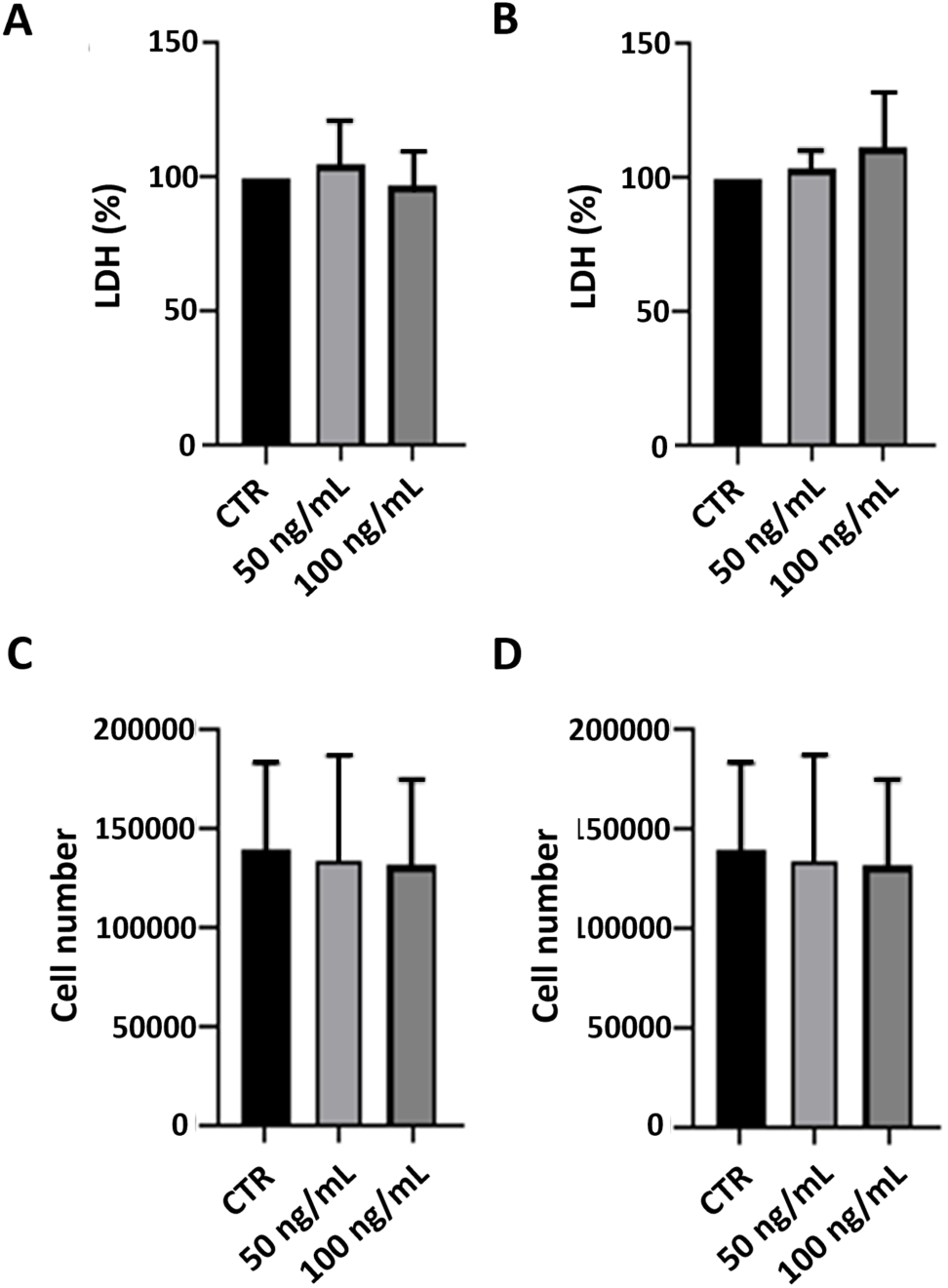
IL-1β had no effect on cell growth or viablity. (A-B) A lactate dehydrogenase assay was used to quantify cell viablity. 24 hours shown in (A) and 72 hours shown in (B). (C-D) Cell counts were used to quantify cell proliferation. 24 hours shown in (C) and 72 hours shown in (D). All data are epressed as mean ± standard deviation of one representative experiment, N=3. One-way ANOVA with a Tukey’s post hoc multiple comparisions test.

We next examined the effect of IL-1β on mitochondrial membrane potential using the cationic dye JC-1. At 24 hours in control cells, mitochondria were well polarized with a branched, tubularl network (Figure 3A). Short term exposure to IL-1β resulted in a loss of membrane polarization. At 72 hours, control cells again had well polarized mitochondria. In contrast to the 24 hour time point however, cells treated with either 50 ng/mL or 100 ng/mL IL-1β regained membrane polarization. The quantification of the ratio of red to green fluorescence confirmed the reduction in polarization at 24 hours (P<0.0001, Figure 3B). After prolonged exposure, the red to green ratio was increased in cells treated with 100 ng/mL IL-1β compared to control cells (Figure P<0.001). Together, these findings show that despite the increase in pro-inflammatory cytokine and ROS production, prolonged culture of IL-1β did not induce any measurable level of cytotoxicty and instead, mitochondrial polarization was restored.

**Figure 3:**
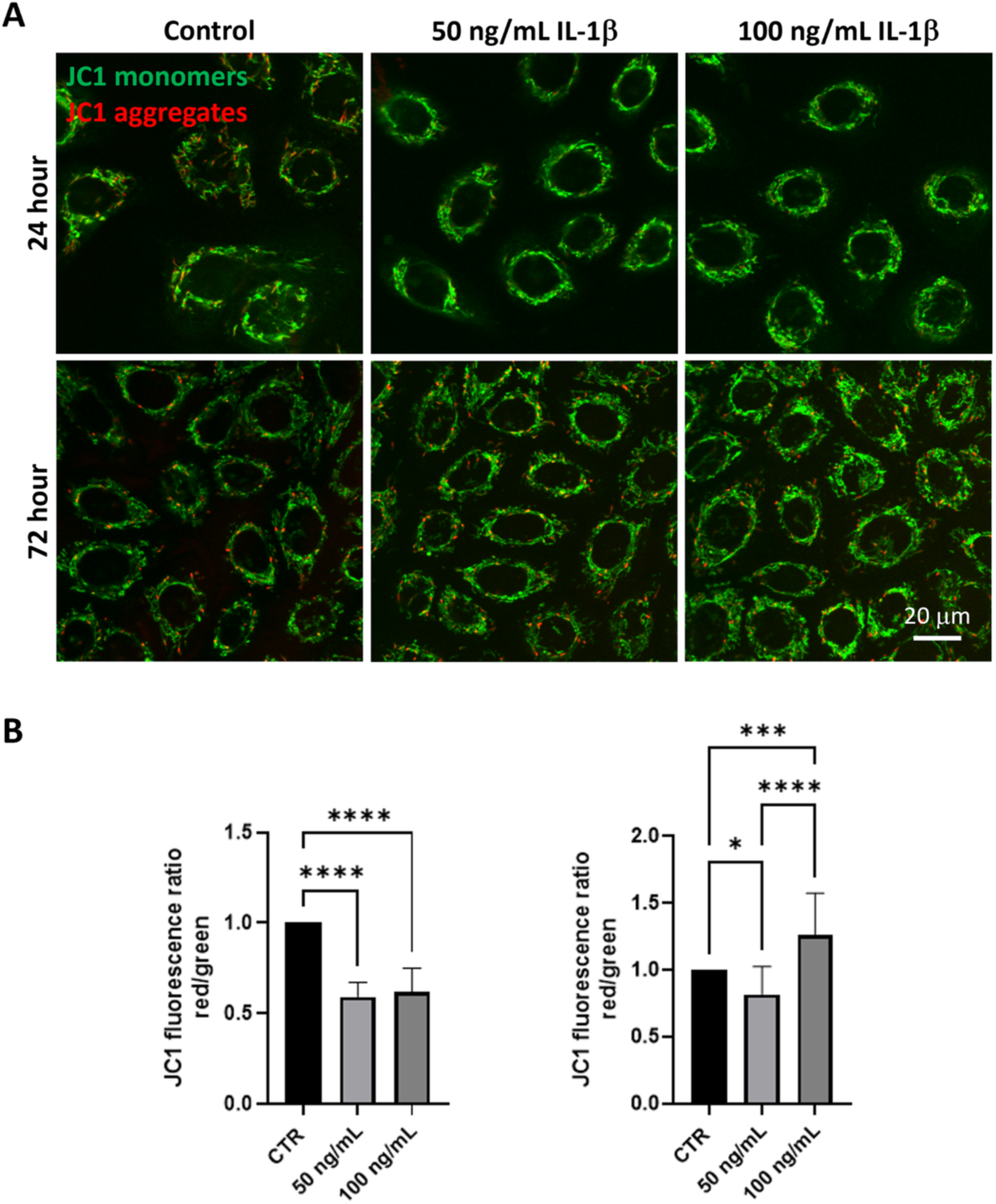
Short term exposure to IL-1β triggered a loss of mitochondrial depolarization that recovered after prolonged exposure in corneal epithelial cells. JC-1 staining was used to quantify mitochondrial polarization. JC-1 monomers indicating depolarized mitochondria are shown in green, JC-1 aggregrates indicating polarized mitochondria are shown in red. (A) JC-1 staining showed a reduction in mitochondrial polarization in after 24 hours of treatment with IL-1β. Mitochondria were smaller and more perinuclear. (B) At 72 hours of treatment, JC-1 aggregate were again visible. Mitochondria exhibited a more elongated, tubular network. (B) Quantification of the red:green fluorescence ratio. There was a decrease in the red:green ratio after treatment with IL-1β that was independent of concentration (p<0.0001). At 72 hours, there was a small reduction in the red:green ratio at 50 ng/mL IL-1β (p=0.013). The red:green ratio was increased following treatment with 100 ng/mL of IL-1β. Scale bar: 20 μm. All data presented as mean ± standard deviation of one representative experiment, N=3. One-way ANOVA with Tukey post hoc multiple comparison test.

### Prolonged exposure to IL-1β increases spare respiratory capacity in corneal epithelial cells

Due to the observed changes in mitochondrial membrane potential, we next examined the effects of IL-1β on cell metabolism. To accomplish this, cells were subject to a Seahorse mitostress assay after prolonged expsoure to either 50 ng/mL or 100 ng/mL IL-1β. Oxygen consumption as a function of time is shown in Figure 4, A-B. For cells cultured in 50 ng/mL IL-1β, there was an increase in total oxygen consumption (basal and non-mitochondrial) without any effect on extracellular acidification (Figure 4, C-D). In contrast to this, there was no increase in total oxygen consumption in cells treated with 100 ng/mL IL-1β; however, there was an increase in extracellular acidification (Figure 4, E-F). ATP-linked respiration and non-mitochondrial oxygen consumption are shown for both concentrations (Figure 4, G-J). There was an increase in ATP-linked respiration in cells treated with 50 ng/mL that coincided with the increase in total OCR (P=0.002). Maximal respiration was increased in cells treated with either concentration of IL-1β (Figure 4, K&M, P=0.019 and P=0.042 for 50 ng/mL and 100 ng/mL, respectively). This resulted in an increase in spare respiratory capacity in treated cells compared to controls, that appeared to increase in a dose-dependent fashion (Figure 4, L&N, P=0.022 and P=0.001). This increase in spare respiratory capacity signals that corneal epithelial cells subject to prolonged IL-1β have increased energy reserves that enhance their ability to respond to additional stress.

**Figure 4:**
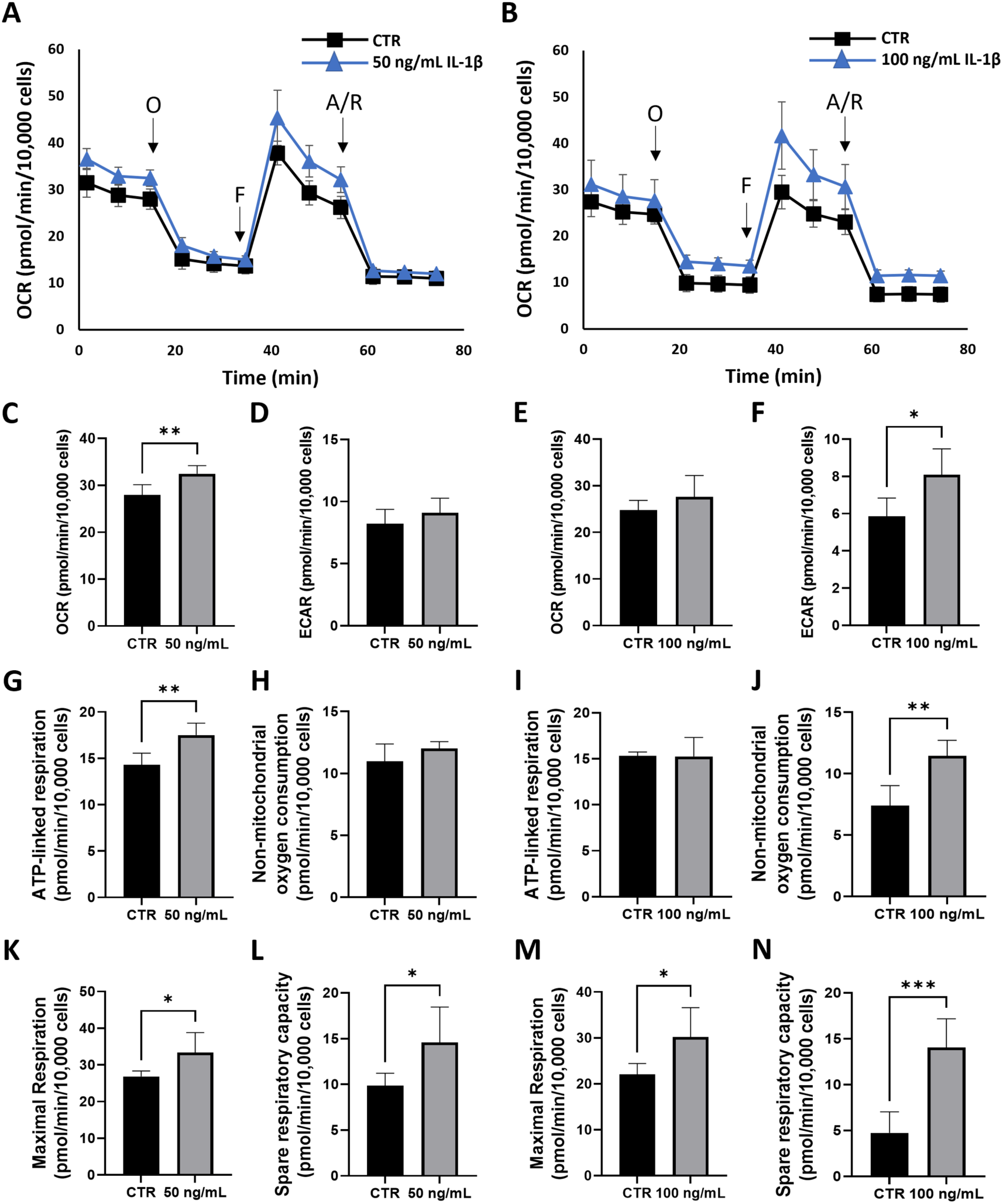
Prolonged exposure to IL-1β increased spare respiratory capacity in corneal epithelial cells. Mitochondrial oxygen consumption was assessed using a MitoStress test. (A-B) Oxygen consumption plotted as a function of time for 50 ng/mL IL-1β (A) and 100 ng/mL IL-1β. (C) 50 ng/mL IL-1β induced a small, but significant increase in total OCR. (D) ECAR was unaffected. (E) 100 ng/mL IL-1β had no affect on total OCR. (F) 100 ng/mL IL-1β increased ECAR. (G) There was an increase in ATP-linked respiration foor 50 ng/mL IL-1β, (H) but no measurable affect on non-mitochondrial oxygen consumption. (I) ATP-linked respiration was unaffected with 100 ng/mL IL-1β, (J) while non-mitochondrial oxygen consumption was increased. (K-N) Both maximal and spare respiratory capacity were increased for 50 ng/mL and 100 ng/mL IL-1β. All data presented as mean ± standard deviation of one representative experiment, N=3. One way ANOVA with Tukeys post hoc multiple comparison test, *p<0.05, **P<0.01, ***p<0.001. OCR: oxygen consumption rate, ECAR: extracelullar acidification rate.

### Prolonged exposure to IL-1β promotes mitochondrial fusion

We next evaluated the effects of IL-1β on PINK1 and BNIP3L/NIX-mediated mitophagy. PINK1 is a mitophagy protein that undergoes degradation by the proteasome in the presence of polarized mitochondria.^33^ Consistent with JC-1 findings showing the restoration of mitochondrial polarization, IL-1β led to a decrease in the expression levels of PINK1 (P<0.05, Figure 5, A-B). We also found a significant reduction in BNIP3L/NIX-mediated mitophagy in the 100 ng/ml treatment group (Figure 5, A&C). To determine the effects of prolonged exposure to IL-1β on mitochondrial structure, we next performed transmission electron microscopy. In control cells, mitochondria appeared normal with the presence of lamellar cristae (Figure 5D). In cells treated with IL-1β, there was evidence of mitochondrial hyperfusion with cristae architeture similar to untreated controls. Mitochondrial fussion is driven, in part, by the need to transfer mitochondrial components during stress to ensure function, as opposed to the onset of mitophagy, which removes damaged mitohcondria.^30,34^ Thus, the observed mitochondrial hyperfusion is consistent with the observed attenuation of mitophagy.

**Figure 5:**
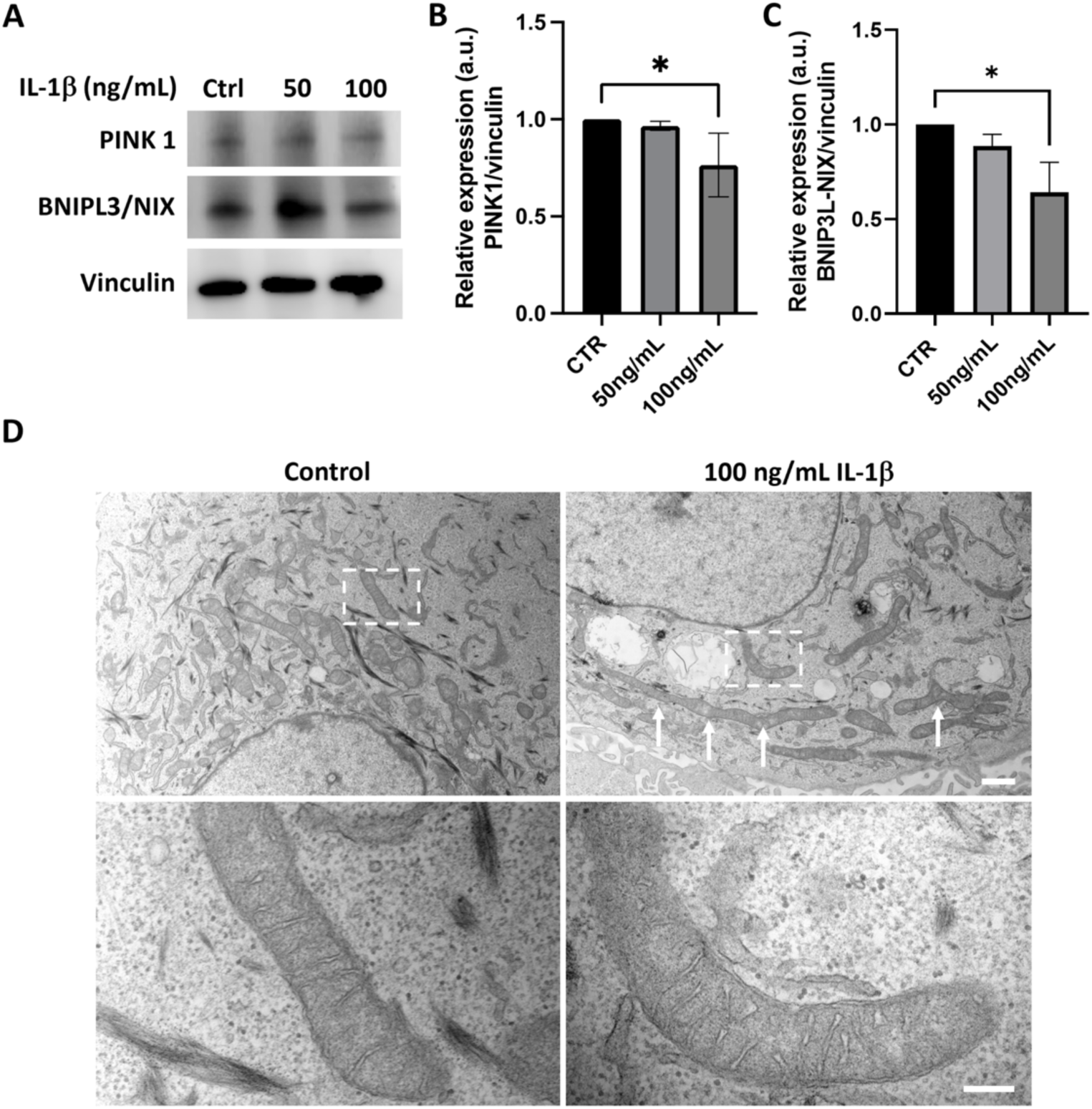
There was a decrease in mitophagy after prolonged exposure to 100 ng/mL IL-1β in corneal epithelial cells. (A-C) Immunoblotting was used to quantify mitophagy proteins, PINK1 and BNIP3L/NIX. Prolonged exposure to IL-1β showed a decrease in both proteins. (B-C) Corresponding densitometry values for (B) PINK1, and (C) BNIP3L/NIX. Proteins were normalized to vinculin. (D) Transmission electron microscopy of cells treated with 100 ng/mL IL-1β for 72 hours showed the presence of intact cristae and mitochondrial hyperfusion (arrows). Bottom panels zoomed region shown by dotted line in the top panels. Zoomed images show simialr cristae morphology between treated and control cells. Scale bar: 1 μm (top panel); scale bar: 200 nm (bottom panel).

### Prolonged exposure to IL-1β induces metabolic rewiring in corneal epithelial cells

To determine the effects of prolonged exposure to IL-1β on cell metabolism, we performed metabolomics using a targeted approach. Cells were again treated with 100 ng/mL IL-1β for 72 hours. The principal component analysis (PCA) showed large differences between control and IL-1β treated cells, with only a small amount of overlap observed between groups (Figure 6A). A supervised analysis by orthogonal projection to latent structures discriminant analysis (OPLS-DA) showed clear ellipse separation, providing further evidence of prominent changes in metabolite composition (Figure 6B). The difference between the PCA and the OPLS-DA is due to the former being an unsupervised method that clusters samples using variance and the latter a supervised method based on regression analysis. We also performed a hierarchiacal cluster analysis illustrated in Figure 6C using dendrograms.

**Figure 6:**
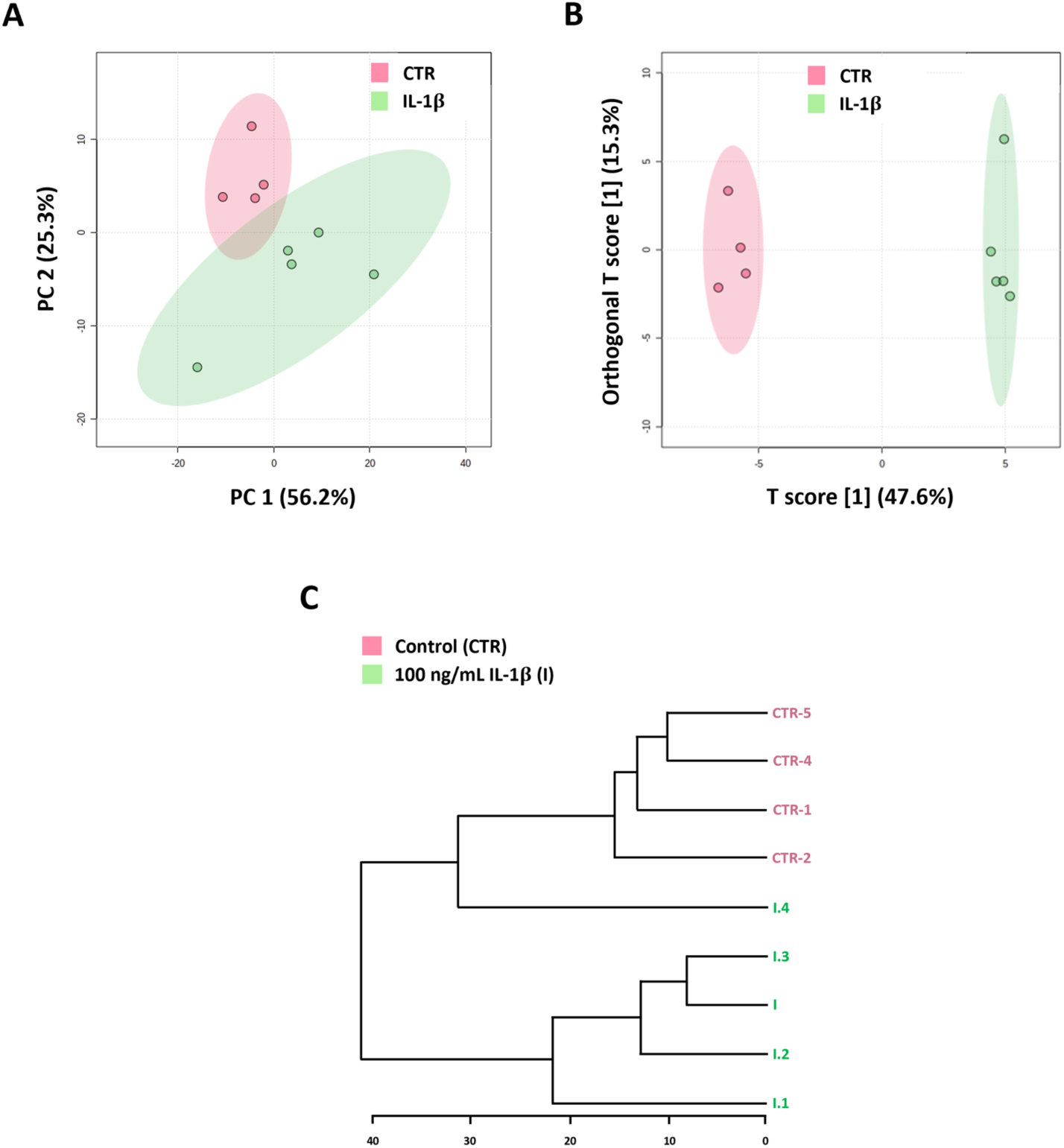
Prolonged exposure to IL-1β induced a shift in the metabolome of corneal epithelial cells. (A) Principal component analysis (PCA) showed distinct differences between control and IL-1β treated cells. (B) Orthogonal projection to latent structures (OPLS-DA) analysis showing complete separation differences in metabolites between groups. (C) Hierarchical cluster analysis dendrogram ilustrating the relationship between controls and cells cultured with IL-1β.

A heat map of differentially expressed metabolites is shown in Figure 7. Some of the most highly expressed metabolites in the control group included glucose-6-phosphate and frutose-6-phosphate, metabolites associated with the first steps of glycolysis. The decrease in these two metabolites in IL-1β treated cells suggests that these metabolites are being consumed due to inceased glycolysis. This would be consistent with the Seahorse assay showing an increase in extracellular acidification, a surrogate marker for glycolysis, after prolonged exposure to 100 ng/mL IL-1β. Seduheptulose 7-phosphase, nicotinamide adenine dinucleotide phosphate (NADP), along with reduced and oxided forms of glutathione, all metabolites involved in the pentose phosphate pathway, were also higher in control cells. The decrease in these metabolites and the corresponding increase in NADPH in cells treated with IL-1β similarly suggests increased flux in the pentose phosphate pathway. In cells treated with IL-1β, increases were also seen in amino acids, and purine and pyrimidine nucleotides. Metoblites related to the TCA cycle, including coenzyme A, nicotinamide adenine dinucleotide, 4-aminobutyrate (GABA), along with fumarate, were also upregulated.

**Figure 7:**
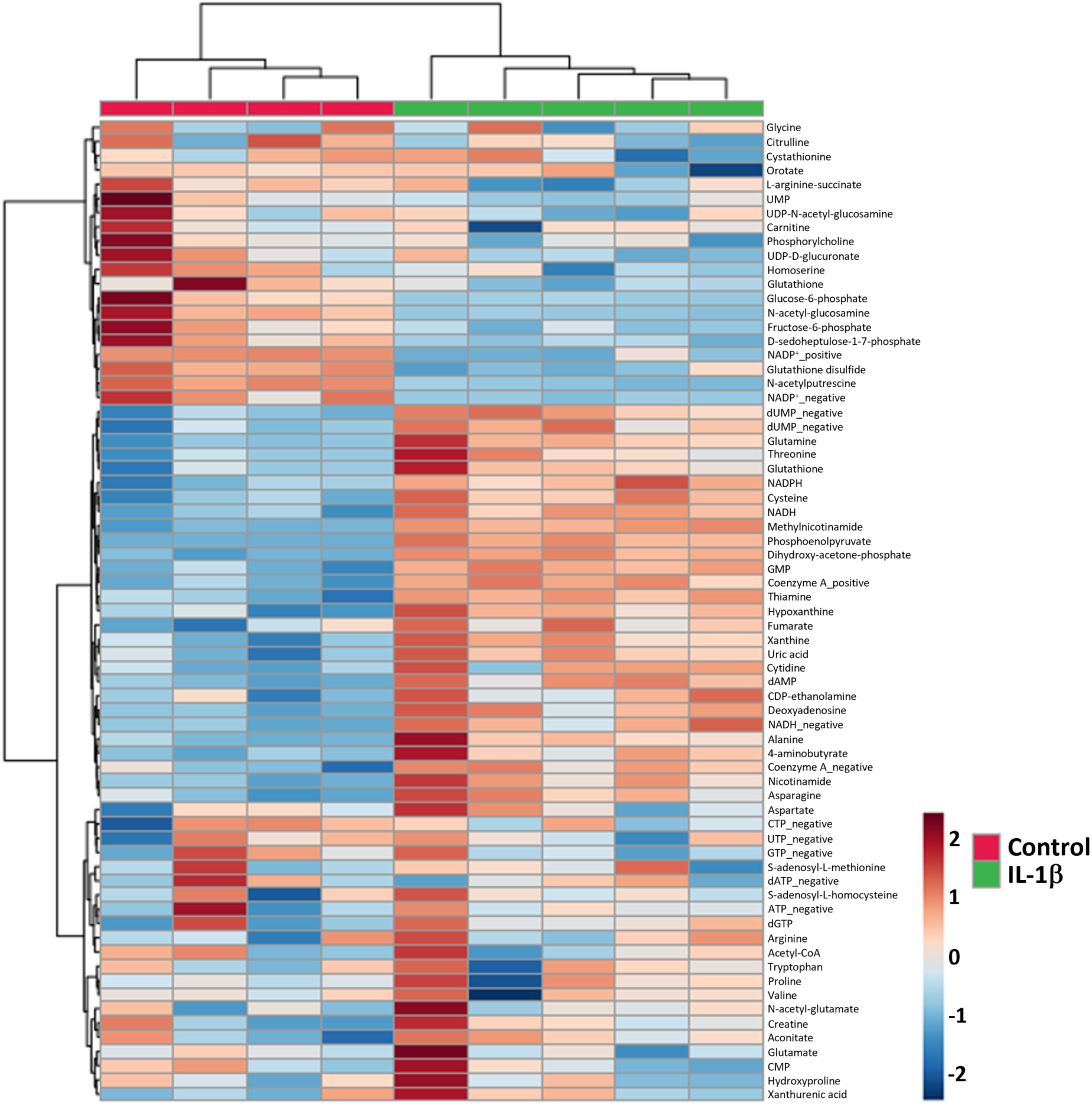
Prolonged exposure to IL-1β induced a shift in the metabolite profile in corneal epithelial cells. A heat map of the differentially expressed metabolites found using a target metabolomics approach. At the top of the heat map, the red color refers to control samples and the green color refers to cells treated in the setting of IL-1β-induced inflammation.

In terms of lipid metabolism, we found an increase in cytidine 5’-diphosphoethanolamine, a metabolite involved in the biosynthesis of ether lipids in IL-1β treated cells. Enrichment analysis of the top 25 metabolites that were increased in response to treatment with IL-1β also showed an upregulation of lipid biosynthetic pathways (Figure 8). The three most enriched pathways were de novo triagcylglycerol biosynthesis, the glycerol-phosphate shuttle, and cardiolipin biosynthesis. Of these, de novo triacylglycerol biosynthesis was the most enriched pathway. Due to the increase in this pathway, we next investigated whether the increase in triacylglycerols were associated with an increase in lipid droplets. Using two different markers for lipid droplets, oil red O and Lipitox, we found an increase in lipid droplets in cells treated with IL-1β (Figure 9, A&B). Quantification of lipid droplets confirmed a significant increase in droplet number using both probes (Figure 9, C&D). These data indicate that corneal epithelial cells respond to inflammatory stress by increasing glycolysis, the TCA cycle, and the pentose phosphate shunt, in addition to changes in lipid homeostasis.

**Figure 8:**
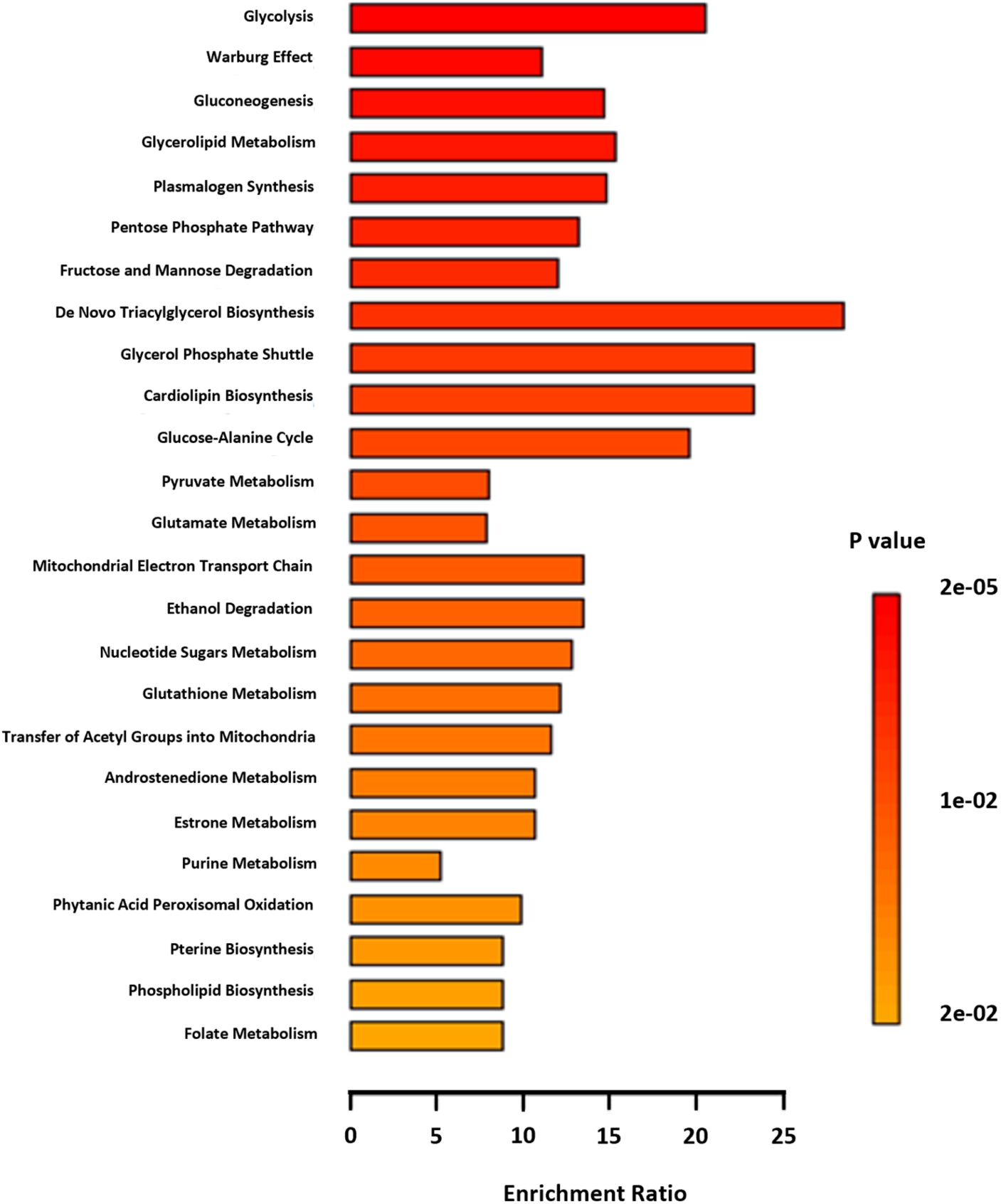
IL-1β increases lipid metabolism in corneal epithelial cells. Enrichment profile for the top 25 upregulated metabolites.

**Figure 9:**
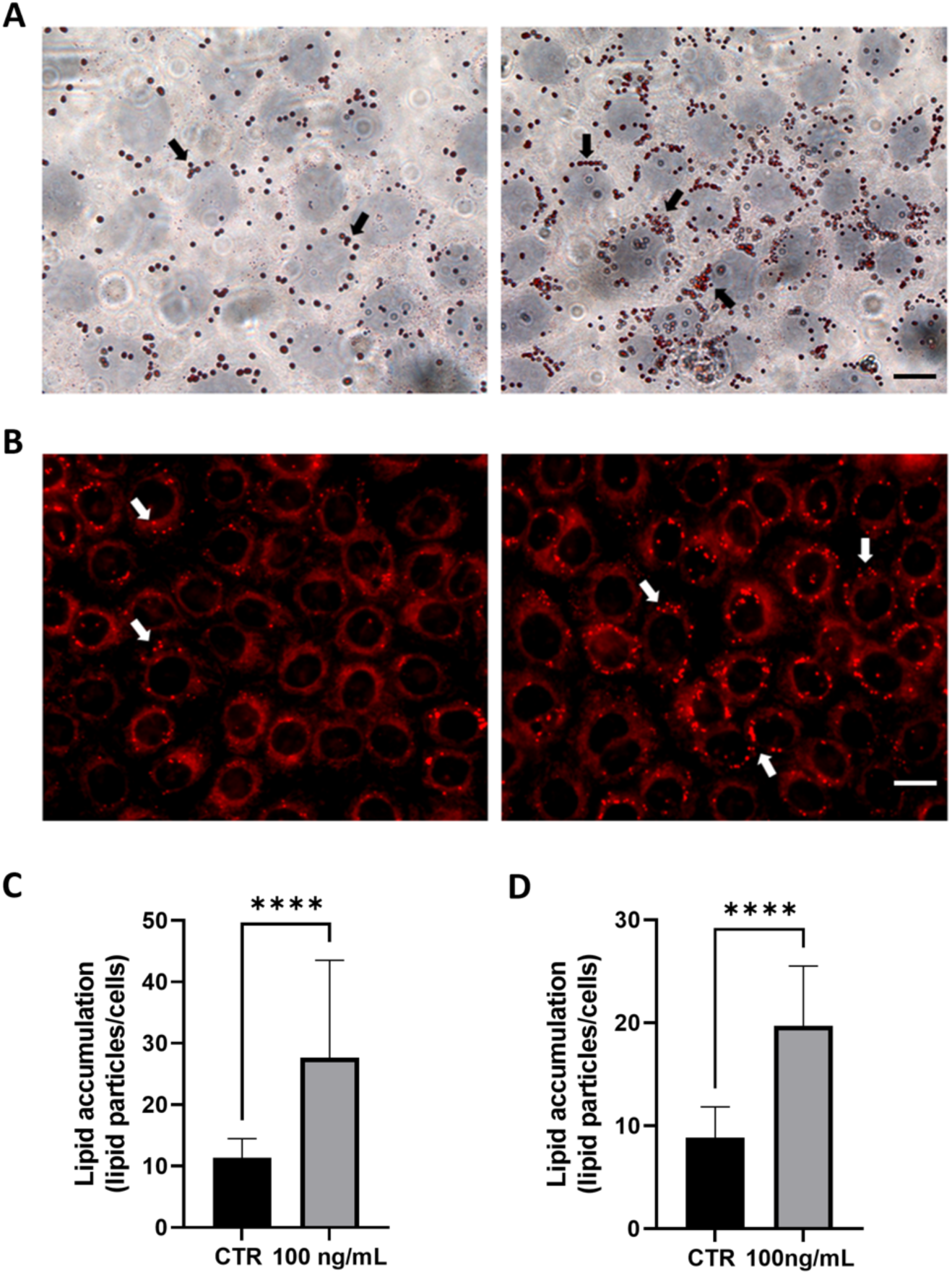
IL-1β induces lipid droplet formation in corneal epithelial cells. (A) Oil red O staining showing lipid droplets present in control cells. There was an increase in lipid droplets in cells treated with 100 ng/mL IL-1β for 72 hours. Black arrows indicate lipid droplets. Scale bar: 50 μm. (B) HCS LipidTox (red) also showed an increase in lipid droplets in cells treated with IL-1β compared to controls. White arrows indicate lipid droplets. Scale bar: 25 μm. (C-D) Lipid droplet quantification for Oil red O (C) and LipidTox (D). Data presented as mean ± standard deviation of 1 representative experiment, N=3. T-test, ****p<0.0001.

## Discussion

In the current study, we investigated the mitochondrial response to IL-1β-mediated inflammatory stress in corneal epithelial cells. We show that IL-1β induced mitochondrial membrane depolarization after short term exposure. This was associated with an increase in pro-inflammatory cytokine production and an increase in ROS. Prolonged expsoure however, led to the paradoxical restoration of mitocondrial membrane potential, despite increased secretion of IL-6 and IL-8 along with a further increase in ROS. The increase in pro-inflammatory cytokines and ROS confirms that IL-1β was not degraded during this time in culture. In terms of mitochondrial function, of the two concentrations tested, 50 ng/ml IL-1β demonstrated a small uptick in basal oxygen consumption without a corresponding change in extracellular acidification. In contrast to this, 100 ng/mL IL-1β triggered an increase extracellular acidification, a surrogate marker of glycolysis. This finding suggests the presence of a fuel shift from oxidative towards glycolytic metabolism at the higher concentration. This finding is consistent with our prior work showing that mild hyperosmolar stress increases oxygen consumption, while it is suppressed at higher levels.^6^

The most striking finding from our metabolic flux experiments however, was the increase is maximum respiratory capacity, that was notably greater at the higher concentration IL-1β that was tested. This finding translated into a higher spare respiratory capacity in the presence of IL-1β. Immunoblotting and ultrastructural analysis confirmed that the increase in spare respiratory capacity was associated with a reduction in PINK1 and BNIP3L/NIX-mediated mitophagy. In healthy, polarized mitochondria, PINK1 is subject to ubiquitin-mediated degradation by the proteasome. Upon membrane depolarization, PINK1 accumulates at the outer mitochondrial membrane to mediate mitophagy.^33^ Thus, the absence of an increase in PINK1 on immunoblotting is consistent with the observed restoration of membrane potential. BNIP3L/NIX-mediated mitophagy, on the other hand, is responsive to physiological stimuli. We have previosuly shown that BNIP3L/NIX expression is highly expressed in the basal layer of the corneal epithelium, decreasing towards the apical surface as cells undergo differentiation.^6^ The mechanism for the decrease in BNIP3L/NIX in response to IL-1β isn’t clear. We speculate that the observed increase in mitochondrial fusion functions to globally downregulate mitophagy as a consequence of decreased fission. Further studies are needed to tease out these findings.

Using a targeted metabolomics approach, we were able to identify specific metabolic changes that occurred during prolonged exposure. The reductions in glucose-6-phosphate, along with an increase in NADPH, suggest an increase in the pentose phosphate pathway. The corneal epithelium is known utilize the pentose phosphate pathway during normal homeostasis. An increase in flux through this pathway would provide both nucleotides and glutathione, the latter of which is needed to buffer the increase in ROS. In addition to the pentose phosphate pathway, there was a clear increase in metabolites involved in glycolysis and the TCA cycle. Importantly, the TCA cycle uses complex II located within the inner mitochondrial membrane to convert succinate to fumarate.^35^ Coinciding with an increase in fumare in IL-1β treated cells was an increase in 4-aminobutyrate, also known as GABA. GABA can readilty be converted to succinate by the enzyme succinate semialdehyde.^36^ Thus, the conversion of succinate to fumarate may account for the increase in spare respiratory capacity in IL-1β treated corneal epithelial cells. Indeed, complex II has been shown to mediate spare respiratory capacity in rat cardiomyocytes.^37^ In marcophages, increased succinate oxidation at complex II has also been shown to enhance mitochondrial membrane potential.^38^ This process leads to an increase in ROS production by driving reverse electron transfer at complex I.

Another potential explanation for the increase in spare respiratory is the increase in cardiolipin that was found. Cardiolipin is a phospholipid that localizes exclusively to the mitochondrial inner membrane.^39^ There cardiolipin interacts with various inner mitochondrial membrane proteins to form the complexes required for the electron transport chain, in addition to supercomplex formation. Cardiolipin also influences the architecture of the inner membrane, where the complexes in the electron transport chain are housed. Hence, the IL-1β mediated increase in cardiolipin may function to increase spare respriatory capacity through inner membrane remodeling. De novo triacylglycerol biosynthesis was also upregulated and was found to be the most enriched pathway. Triacylglyerols are neutral lipids that are formed in the endoplasmic reticulum (ER). The accumulation of triacylglycerols in the ER leads to the biogenesis of lipid droplets that are released into the cytoplasm.^40^ Lipid droplets function as storage depots for lipids until they are needed for energy and to protect against lipotixicity.^41^ In addition, lipid droplets are also now well recognized as mediators of signal transduction and interact directly with mitochondria to maintain organelle homeostasis and promote the removal of damaged lipids during cell stress.^40–43^ We were able to quantify lipid droplets using two different probes and confirmed an increase in lipid droplet formation. In our unpublished work, we have als observed the formation of lipid droplets in response to hyperosmolar and dessicating stress. While little is known about the function of lipid droplets in corneal epithelial cells, this finding represents an exciting avenue for future studies.

## Conclusion

In summary, the potential for prolonged exposure of IL-1β to trigger metabolic adaptation in corneal epithelial cells has important implications for the ocular surface and the concept of immune tone. Metabolic adaptation is a survival response that functions to prime the cornea, making it more resilient against envinromental insults, trauma, and even infectious pathogens during contact lens wear. The findings presented here pave the way for further work to determine whether inflamation is truly beneficial to the ocular surface, and if so, when does inflammation shift from beneficial to pathological.

## Acknowledgments

None.

## Disclosure

The author(s) report no conflicts of interest in this work.

## Notes

### Competing Interest Statement

The authors have declared no competing interest.

